# Comparative genomic insights into the evolution of *Halobacteria*-associated “*Candidatus* Nanohaloarchaeota”

**DOI:** 10.1101/2022.05.20.492899

**Authors:** Dahe Zhao, Shengjie Zhang, Sumit Kumar, Heng Zhou, Qiong Xue, Wurunze Sun, Jian Zhou, Hua Xiang

## Abstract

The phylum “*Candidatus* Nanohaloarchaeota” is a representative halophilic lineage within DPANN superphylum. They are characterized by their nanosized cells and symbiotic lifestyle with *Halobacteria*. However, the development of the symbiosis remains unclear for the lack of genomes located at the transition stage. Here, we performed a comparative genomic analysis of “*Ca*. Nanohaloarchaeota”. We propose a novel family “*Candidatus* Nanoanaerosalinaceae” represented by five de-replicated metagenome-assembled genomes obtained from hypersaline sediments and the enrichment cultures of soda-saline lakes. Phylogeny analysis reveals that the novel family are placed at the root of the family “*Candidatus* Nanosalinaceae” including the well-researched taxa. Most members of “*Ca*. Nanoanaerosalinaceae” contain lower proportion of putative horizontal gene transfers from *Halobacteria* than “*Ca*. Nanosalinaceae”, while they maintain moderately acidic proteomes for hypersaline adaptation of “salt-in” strategy, suggesting that “*Ca*. Nanoanaerosalinaceae” have not established an intimate association with *Halobacteria*, and may descend from an intermediate stage. Functional prediction discloses that they exhibit divergent potentials in carbohydrate and organic acids metabolism, and environmental responses. Historical events reconstruction illustrates that the involved genes acquired at the putative ancestors possibly drive the evolutionary and symbiotic divergences. Globally, this research on the new family “*Ca*. Nanoanaerosalinaceae” enriches the taxonomic and functional diversity of “*Ca*. Nanohaloarchaeota”, and provides insights into the evolutionary process of “*Ca*. Nanohaloarchaeota” and their *Halobacteria*-associated symbiosis.

**Importance:** DPANN superphylum is a group of archaea widely distributing in various habitats. They generally have small cells, and perform a symbiotic lifestyle with other archaea. The archaeal symbiotic interaction is important to understand microbial community. However, the formation and evolution of the symbiosis between the DPANN lineages and other diverse archaea remain unclear. Based on phylogeny, hypersaline adaptation, functional potentials, and historical events of “*Ca*. Nanohaloarchaeota”, a representative phylum within the DPANN superphylum, we report a novel family descending from an intermediate stage, and we illustrate the evolutionary process of “*Ca*. Nanohaloarchaeota” and their *Halobacteria*-associated symbiosis. Furthermore, we find the acquired genes involved in carbohydrate and organic acids metabolism and environmental responses possibly drive the evolutionary and symbiotic divergences. Altogether, this research helps in understanding the evolution of the archaeal symbiosis, and provides a model for the evolution of the other DPANN lineages.

## Introduction

DPANN superphylum (an acronym of five candidate phyla names, “*Candidatus* Diapherotrites”, “*Candidatus* Parvarchaeota”, “*Candidatus* Aenigmarchaeota”, “*Candidatus* Nanoarchaeota”, and “*Candidatus* Nanohaloarchaeota”) is a group of archaea with nanosized cell and small genome size (1–3). Despite different classifications amid Genome Taxonomy Database (GTDB) and NCBI taxonomy database (4), more and more lineages are classified into the DPANN superphylum, including “*Candidatus* Micrarchaeota” (5), “*Candidatus* Woesearchaeota” (6), “*Candidatus* Pacearchaeota” (6), “*Candidatus* Huberarchaeota” (7), and “*Candidatus* Undinarchaeota” (8). The DPANN group formed at the early stage of life’s evolution (8–12). A general symbiotic lifestyle is proposed from the reduced metabolic potentials of most members (3, 5, 6, 13). The symbiosis was demonstrated based on the cocultures of DPANN lineages and their hosts (2, 14–18). Many lineages, like “*Ca*. Nanoarchaeota”, “*Ca*. Huberarchaeota”, “*Ca*. Nanohaloarchaeota” and “*Ca*. Aenigmarchaeota” were predicted to exchange genes with their respective hosts via horizontal gene transfer (HGT) (8, 18–21). However, it remains unknown how the DPANN lineages form symbionts with diverse taxa.

“*Ca*. Nanohaloarchaeota” was one of the first five phyla in the DPANN group (1). This phylum is widely distributed in the (hyper)saline habitats (3, 22–26) by harnessing the energetically favorable “salt-in” strategy (22, 24) like their host *Halobacteria* (27). “*Ca*. Nanohaloarchaeota” cells were revealed to have a tightly symbiotic relationship with the class *Halobacteria* as demonstrated by their cocultures (16, 17). In some “*Ca*. Nanohaloarchaeota” genomes, long “SPEARE” proteins containing serine protease, adhesion, and restriction endonuclease domains were supposed to function in attachment and invasion of hosts (16). Remarkably, nanohaloarchaeon “*Candidatus* Nanohalobium constans” LC1Nh exhibits mutualistic symbiosis with the host under the conditions with glycogen or starch as a carbon source (17). “*Ca*. Nanohaloarchaeota” shared 21% of sisterhood relationships with *Halobacteria* (8), and similar HGT events were also reported in other works of literature (21). However, because of the protein adaptation to high salinity in the cytoplasm, some of the close relationships may be the result of compositional biases from convergent evolution (8, 22, 24). Based on the cocultivation and genomic prediction, “*Ca*. Nanohaloarchaeota” were considered aerotolerant anaerobes with a lifestyle of sugar fermentation, while the hosts generally perform aerobic respiration (3, 17, 22, 23). In addition, all these reports on “*Ca*. Nanohaloarchaeota” were focused on one family, i.e., “*Candidatus* Nanosalinaceae” (see the Results and Discussion below).

In this research, we report five de-replicated metagenome-assembled genomes (MAGs) of a novel family named “*Candidatus* Nanoanaerosalinaceae” in “*Ca*. Nanohaloarchaeota”. They were obtained from hypersaline sediments or the enrichment culture of a soda-saline lake. We find that this family forms a separate clade in the phylogenetic tree, and some members contain a lower proportion of horizontally acquired genes from *Halobacteria*. These results indicate that this family emerged at an early stage. Furthermore, we performed the amino acid composition, functional gene prediction, and comparative genome analyses to illustrate the evolutionary process of “*Ca*. Nanohaloarchaeota”. Based on these findings, we infer the formation and evolution of symbiosis with *Halobacteria*.

## Results and Discussion

### Acquisition of the genomes of the novel family “*Ca*. Nanoanaerosalinaceae”

To obtain the genomes of novel taxa, we performed enrichment culture. Briefly, five deep sediment samples were used for enrichment culture with the addition of inorganic salts, nutrients, and antibiotics (Table S1). The deep sediment samples have been described in our previous research (28). After incubation of about 210 days, the microbial cells (with insoluble matter) were collected using centrifugation, and then the metagenome sequencing and genome binning were performed. The details are described in Methods. Additionally, from eighteen metagenomes of brine and surface sediment samples provided in our earlier research (23), genome binning was re-assembled by means of more tools in this research. After de-replication, taxonomy annotation (based on GTDB), and quality estimation (more than 75% completeness and less than 5% contamination), a total of 10 metagenome-assembled genomes (MAGs) affiliated with “*Ca*. Nanohaloarchaeota” were obtained (Table 1). In the description below, we will mainly follow the GTDB taxonomy if not specified. To systematically decipher the phylogenetic analyses and comparative genomics of the “*Ca*. Nanohaloarchaeota”, we collected 235 genomes affiliated with “*Ca*. Nanohaloarchaeota” or its closely related phyla (“*Ca*. Aenigmarchaeota”, EX4484-52, PWEA01, and QMZS01; NATU short for all five phyla). In total, there were 116 NATU genomes with completeness of more than 75% and contamination of less than 5%, including 24 “*Ca*. Nanohaloarchaeota” genomes (Table S2). Meanwhile, the firstly reported genome J07AB56 (3) and one complete genome assembled from the metagenome of Nha-CHl enrichment culture (16) were retained for the following research. The taxonomic annotation of GTDB-Tk pipeline reveals that all of the 26 “*Ca*. Nanohaloarchaeota” genomes belong to the order “*Candidatus* Nanosalinales”, of which 19 are affiliated with the family “*Ca*. Nanosalinaceae”, and the other 7 are not classified (including the family “*Ca*. Nanoanaerosalinaceae”; described below).

**Table 1.** General genomic features of the “*Ca*. Nanohaloarchaeota” metagenome-assembled genomes (MAGs) obtained in this research

| MAG | WGS accession number | Size (bp) | Contigs number | N50 (bp) | GC (mol%) | Comp (%) | Cont (%) | Taxonomy at family level | Source | Salinity (%)<sup>b</sup> |
| --- | --- | --- | --- | --- | --- | --- | --- | --- | --- | --- |
| NHA20 | JALDAE000000000 | 840,242 | 60 | 21,270 | 41.20 | 79.91 | 0.00 | Nanoanaerosalinaceae | Sediment | 32 |
| NHA21 | JALDAF000000000 | 1,130,266 | 78 | 19,632 | 56.09 | 78.94 | 0.00 | Nanoanaerosalinaceae | Enrichment (DK15A)<sup>a</sup> | 15 |
| NHA22 | JALDAG000000000 | 948,638 | 56 | 24,269 | 38.81 | 80.37 | 0.47 | Nanoanaerosalinaceae | Sediment | 20.7 |
| NHA23 | JALDAH000000000 | 849,024 | 82 | 17,982 | 38.05 | 78.50 | 0.93 | Nanoanaerosalinaceae | Sediment | 27 |
| NHA24 | JALDAI000000000 | 762,957 | 106 | 9,734 | 38.12 | 76.53 | 2.80 | Nanoanaerosalinaceae | Sediment | 27 |
| NHA25 | JALDAJ000000000 | 1,027,595 | 1 | 1,027,595 | 48.48 | 86.14 | 0.00 | Nanosalinaceae | Enrichment (DK24A)<sup>a</sup> | 24 |
| NHA26 | JALDAK000000000 | 899,074 | 37 | 74,505 | 40.69 | 82.36 | 2.80 | Nanosalinaceae | Brine | 32 |
| NHA27 | JALDAL000000000 | 847,177 | 240 | 4,323 | 40.05 | 77.65 | 4.35 | Nanosalinaceae | Sediment | 27 |
| NHA28 | JALDAM000000000 | 844,933 | 64 | 20,433 | 39.87 | 82.20 | 1.40 | Nanosalinaceae | Sediment | 22.4 |
| NHA29 | JALDAN000000000 | 886,350 | 48 | 24,696 | 44.82 | 84.27 | 0.93 | Nanosalinaceae | Brine | 27.3 |

Note: a, the enrichment is under the anaerobic condition; "A" means sodium acetate was added; details are provided in Table S1. b, the salinity is for the pond or the enrichment media. Comp, Completeness; Cont, Contamination.

### Phylogeny of the family “*Ca*. Nanoanaerosalinaceae” within the phylum “*Ca*. Nanohaloarchaeota”

In the phylogenomic tree based on 122 single-copy conserved proteins, ribosome proteins, and the 16S rRNA gene (Figs. S1–S3, and Tables S3, S4), NATU are located in the DPANN group, while the class *Halobacteria* (the host of “*Ca*. Nanohaloarchaeota”) are affiliated with the phylum *Halobacteriota*, whose subordinates belong to the phylum *Euryarchaeota* of the classical taxonomy (4). The phyla “*Ca*. Nanohaloarchaeota” and EX4484-52 cluster together (Figs. S4–S6). There seem to be three family-level lineages with bootstrap supports of more than 75%, including “*Ca*. Nanosalinaceae”, “*Ca*. Nanoanaerosalinaceae” and AB_1215_Bin_137 (Figs. 1, S4–S6). In fact, AB_1215_Bin_137 is a MAG obtained from one of the Guaymas Basin (Gulf of California) sediment samples (29). It is classified into the order “*Ca*. Nanosalinales” by GTDB-Tk (Table S2), but it shares a relatively long distance with the other two families (Figs. 1, S4, S5, S6), and its 16S rRNA gene shares similarities of less than 82.0% (order threshold) (30) with them (Fig. S7). Therefore, it may represent a novel order. “*Ca*. Nanoanaerosalinaceae” is placed at the root of the phylogenetic trees (Figs. 1, S4–S6). The AAI values of the predicted proteome between the genomes of the families “*Ca*. Nanosalinaceae” and “*Ca*. Nanoanaerosalinaceae” are generally less than 45%, the family boundary according to the previous literature (31); and those among the genomes of the same family are generally greater than 45% (Fig. 1). The 16S rRNA gene identity analysis backs the classification of “*Ca*. Nanoanaerosalinaceae” (Fig. S7) according to the taxonomic thresholds of sequence identities for order and family (82.0% and 86.5%, respectively) (30). In conclusion, we propose a new family placed at the root of the “*Ca*. Nanohaloarchaeota” phylogenies. Five genomes provided in this research were obtained from sediment samples or anaerobic enrichment (Table 1), while NHA-2 was acquired from the sediment sample of a pond with a salinity of 20% in the previous research (23). Considering their habitats, we named them “*Ca*. Nanoanaerosalinaceae”. Markedly, the taxonomy is different from the previous report (17), in which the taxa of the family “*Ca*. Nanosalinaceae” was classified into three classes. The possible reason may be that the other two lineages, AB_1215_Bin_137, and the family “*Ca*. Nanoanaerosalinaceae” placed at the root of “*Ca*. Nanohaloarchaeota” were not included in the phylogenomic analysis. Moreover, the well-researched “*Ca*. Nanohaloarchaeota” taxa belong to the family “*Ca*. Nanosalinaceae”, including the two cocultures with *Halobacteria* (16, 17). According to 95% AAI and ANI for species boundaries (31), six replicated genomes with low completeness or high contamination were removed (Fig. 1, Table S5). During downloading the genomes, we found that 10 assemblies were classified in the “*Ca*. Nanohaloarchaeota” (under taxid 1462430) of DPANN group, while 29 were in the class “*Ca*. Nanohaloarchaea” (under taxid 1051663) of *Euryarchaeota* in NCBI taxonomy database. However, it is clear that they share a very close relationship with each other. The misinterpretation was considered as the result of inadequate outgroup representation (1).

**Fig. 1.**
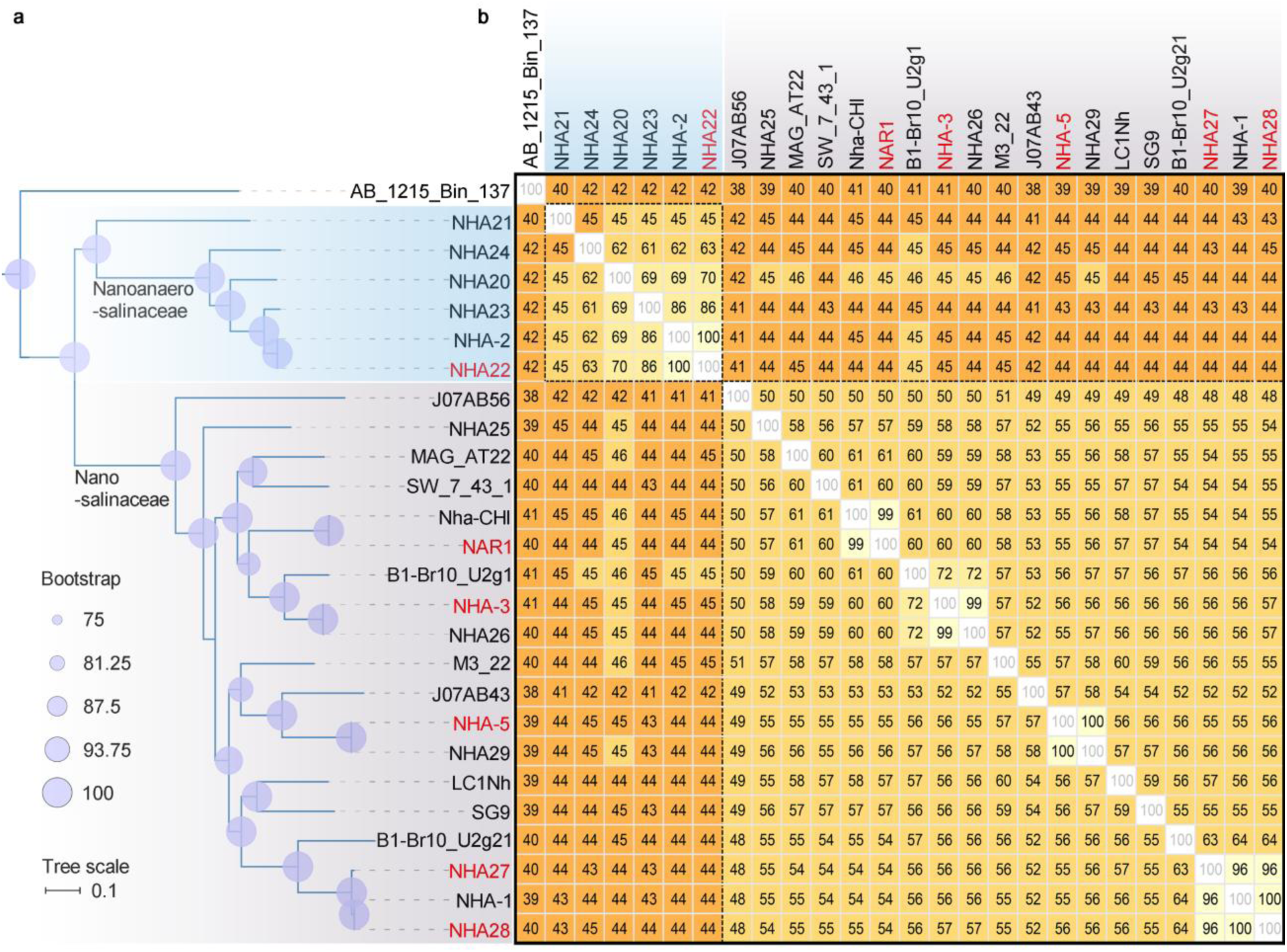
Phylogeny of the phylum “*Ca*. Nanohaloarchaeota”. (a) Phylogenomic trees based on the 122 single-copy ubiquitous proteins in GTDB. It is obtained by pruning the tree in Fig. S4. Briefly, the best-fit model of LG+F+G4 is chosen, and a consensus tree based on ultrafast bootstrap approximation of 1,000 times is presented. (b) Average amino acid identity (AAI) matrix among the genomes of “*Ca*. Nanohaloarchaeota”. The data are rounded by omitting decimal fractions smaller than 0.5 and counting all others (including 0.5) as 1. The background (from yellow to orange) is painted according to the threshold AAIs of species, genus, and family (95.0, 65.0, and 45.0%, respectively). The genomes sharing an AAI of more than 95.0% with other genome of higher completeness or low contamination are marked by red, and they are abandoned in the following research.

### “Ca. Nanoanaerosalinaceae” emerged at an intermediate stage of Halobacteria-associated symbiosis

Considering the genes exchange between “*Ca*. Nanohaloarchaeota” and *Halobacteria* as a result of their intimate association with each other in natural habitats (8), we detected the putative horizontal gene transfers (HGTs) using the HGTector tool, which is founded on sequence homology search hit distribution statistics (32). First, we created a GTDB archaeal taxonomy-based database (described in Methods) because the default database follows NCBI taxonomy, in which 29 genomes were incorrectly classified into the class “*Ca*. Nanohaloarchaea” (taxid 1051663) of *Euryarchaeota*. Afterward, the HGT events were predicted in the 14 representative genomes of the family “*Ca*. Nanosalinaceae” by following the tutorials. The result shows that HGT events could be found in them (Table S6), and these genomes harbor high proportions (46.25–79.25%) of horizontally acquired genes from *Halobacteria* (Fig. 2). Although the tools are different, a similar conclusion could be drawn as the previous report (8).

**Fig. 2.**
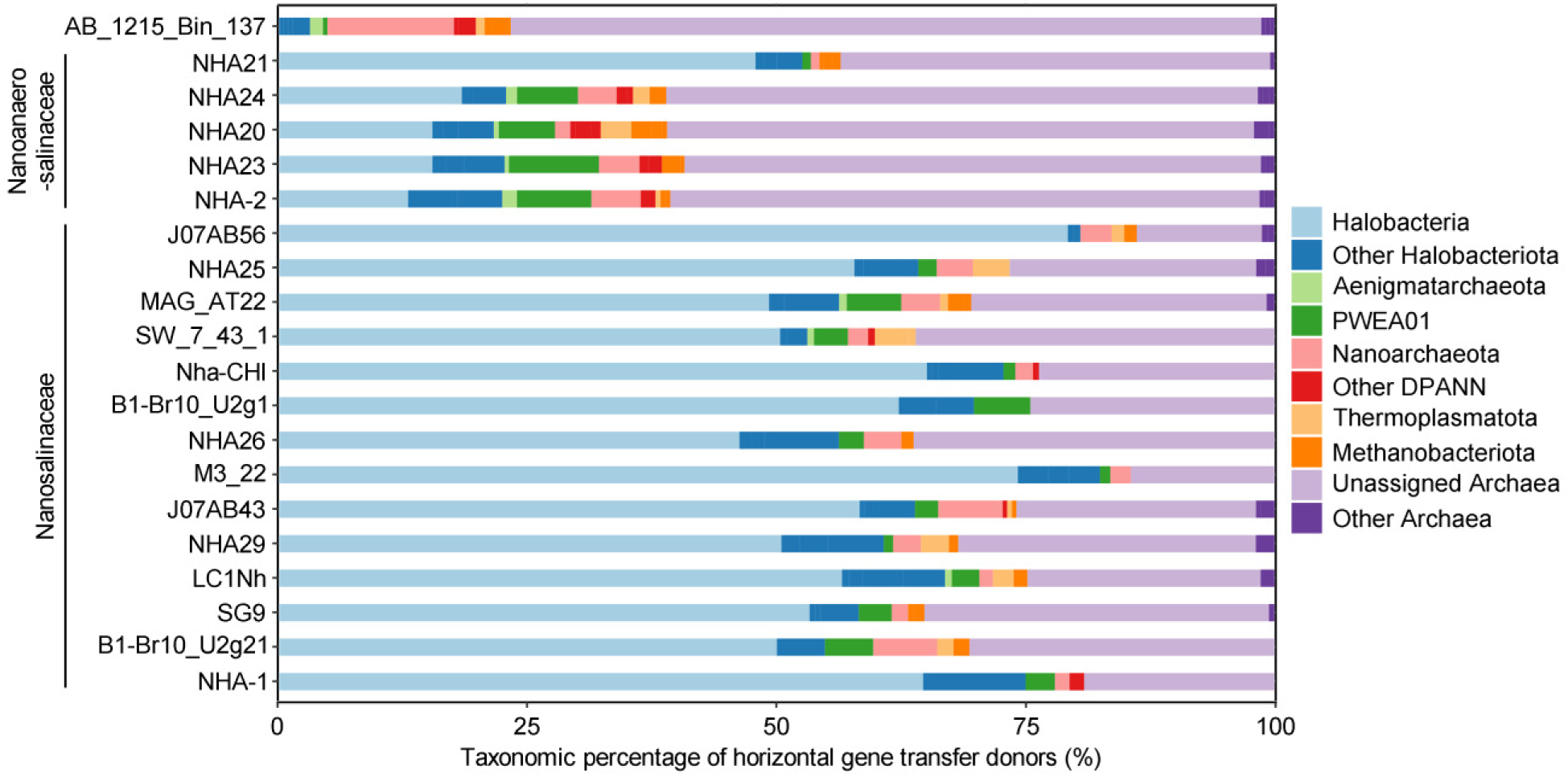
Horizontal gene transfer inference in the genomes of the phylum “*Ca*. Nanohaloarchaeota”. The taxonomic percentage of horizontal gene transfer donors for each MAG/SAG is inferred using the HGTector pipeline with a GTDB taxonomy-based database based on the archaeal genomes in GTDB Release 202. The genomes that are affiliated with the family “*Ca*. Nanosalinaceae” and “*Ca*. Nanoanaerosalinaceae” are marked.

Furthermore, the HGTector tool was used for HGTs prediction of other lineages. The result exhibited that among the 228 HGT events in AB_1215_Bin_137, even none was detected with *Halobacteria* as donors (Fig. 2, Table S6). Combining the phylogeny, AB_1215_Bin_137 lineage might emerge before its ancestor meets *Halobacteria*, while “*Ca*. Nanosalinaceae” may form an intimate association with *Halobacteria*. In “*Ca*. Nanoanaerosalinaceae”, only 12.94–18.33% of the HGT events are found in NHA24, NHA20, NHA23, and NHA-2, while 47.86% in NHA21 (Fig. 2). In other words, “*Ca*. Nanoanaerosalinaceae” members commonly harbor moderate proportions of horizontally acquired genes from *Halobacteria*, and they possibly arise at an intermediate stage. Notably, NHA21 harbors a similarly high proportion of horizontally acquired genes from *Halobacteria* compared to “*Ca*. Nanosalinaceae”. However, it is located at the “*Ca*. Nanoanaerosalinaceae” with almost 100% bootstrap supports in the phylogenetic trees (Figs. 1, S5, S6), and its proteome shares higher similarities with the other members of “*Ca*. Nanoanaerosalinaceae” than those of “*Ca*. Nanosalinaceae” (Fig. 1). Therefore, simultaneously considering its different amino acid composition of proteome and functional profile (described below), NHA21 is assumed to evolve through another path different from the other four species of “*Ca*. Nanoanaerosalinaceae”. In addition, we could observe that the main donors of horizontally acquired genes in most “*Ca*. Nanoanaerosalinaceae” genomes were predicted as unassigned archaea over *Halobacteria* (Fig. 2). It suggests that they may not build an intimate connection with *Halobacteria* or have a wide range of archaeal hosts.

### Hypersaline adaptation of “*Ca*. Nanoanaerosalinaceae” provide new indicators for the evolution

It is widely believed that “*Ca*. Nanosalinaceae” members adopt the “salt-in” strategy to resist the high osmotic pressure of hypersaline environments (22, 24), while AB_1215_Bin_137 inhabiting deep-sea sediment (29) may not be obligate to adapt to the extreme osmotic pressure. During the evolution of the “salt-in” strategy, the isoelectric point profiles of the predicted proteome became acid-shifted (22, 24), because the negatively charged amino acids can maintain the stability and activity of proteins in the hypersaline conditions (33). Consequently, we compared the isoelectric point profiles and amino acid compositions of the predicted proteomes between “*Ca*. Nanoanaerosalinaceae” and reference lineages. Our results support that the isoelectric point profiles of the predicted proteomes of the family “*Ca*. Nanosalinaceae” were acid-shifted (Fig. S8), and their average isoelectric points range from 4.87–5.59 (Table S7). They are close to the three *Halobacteria* references (4.71–4.83). Notably, AB_1215_Bin_137 that does not adopt the “salt-in” strategy displays an alkali-shifted isoelectric point profile and an average isoelectric point of 8.10 (Fig. 3a, Table S7). In “*Ca*. Nanoanaerosalinaceae”, NHA21 displays a similar isoelectric points profile as LC1Nh, one representative strain of “*Ca*. Nanosalinaceae” (Fig. 3a). Correspondingly, its average isoelectric point is low to 5.04 (Table S7). The isoelectric points profiles of NHA24, NHA20, NHA23, and NHA-2 are also acidic, but the acid-shift is weak (Fig. 3a). Their average isoelectric points range from 5.69–6.16 (Table S7), a little higher than that of “*Ca*. Nanosalinaceae” and NHA21. It suggests that “*Ca*. Nanoanaerosalinaceae” may maintain moderately acidic proteomes and concentration of intracellular inorganic salt.

**Fig. 3.**
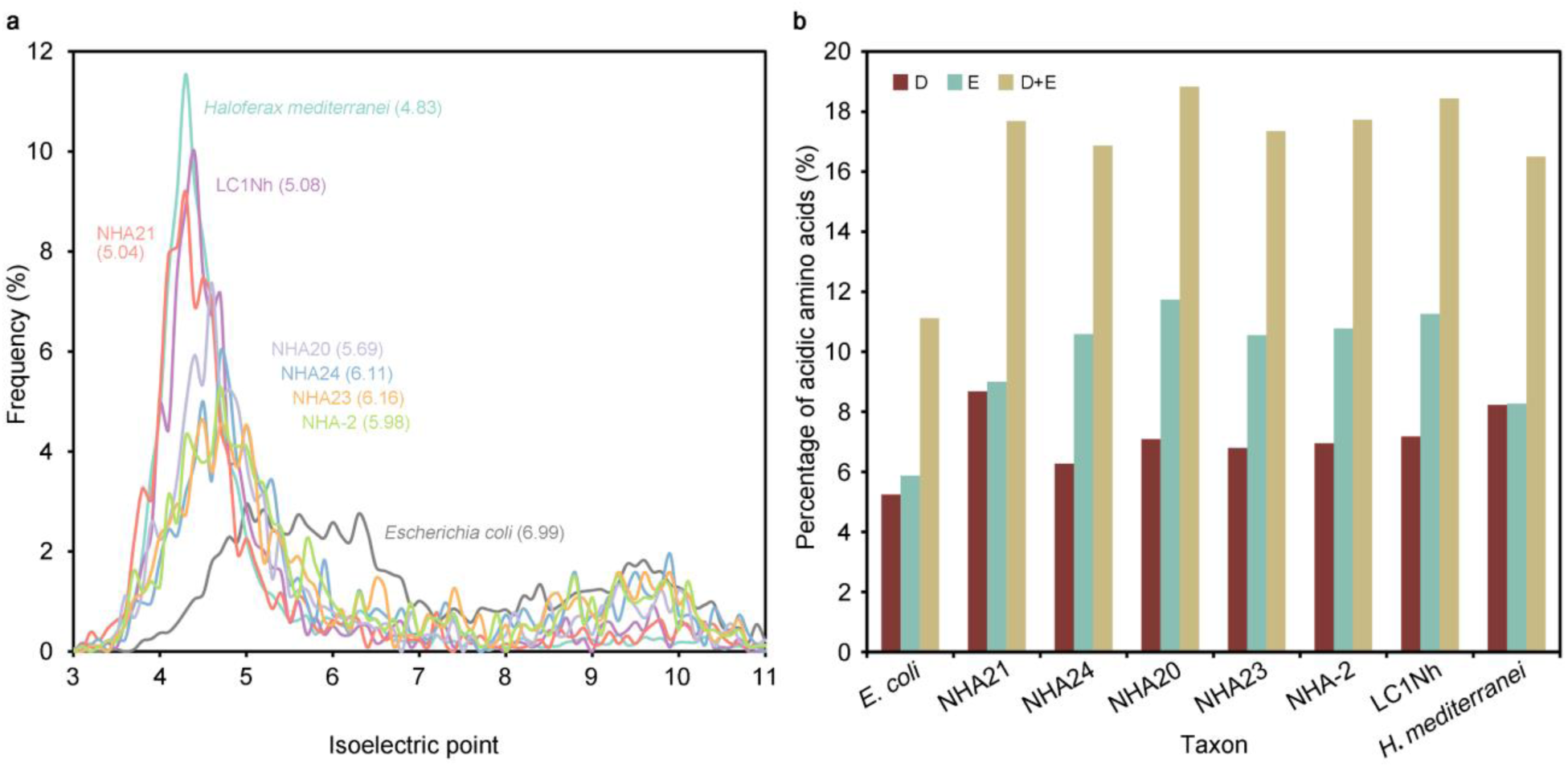
Comparison of isoelectric point profiles and amino acids compositions. (a) Isoelectric point profiles of the predicted proteomes of “*Ca*. Nanoanaerosalinaceae” and reference species. Horizontal coordinates are isoelectric points, and longitudinalcoordinates are the frequencies of proteins in the proteomes at each isoelectric point. The isoelectric point of each protein is predicted based on the amino acid sequence. The isoelectric point profiles of bin width 0.1 are shown. *Haloferax mediterranei* ATCC 33500 (GCA_000306765.2) and *Escherichia coli* O157:H7 str. Sakai (GCA_000008865.2) were shown as acid-shifted “salt-in” halophile and non-halophile. The number indicates the average isoelectric point in the round brackets, and more details are presented in Table S7. (b) Percentage of acidic amino acid glutamate, aspartate, and the sum. The composition is calculated from the predicted proteome based on the genome sequence. D, aspartate; E, glutamate; D+E, the sum of glutamate and aspartate.

Considering such an acidic proteome of NHA21, we further compared the amino acid composition, especially the acidic amino acids glutamate and aspartate. In comparison with the non-halophiles like *Escherichia coli*, the sum of the mole percentage of glutamate and aspartate is about 11%, while those of the “salt-in” *Halobacteria*, “*Ca*. Nanosalinaceae”, and “*Ca*. Nanoanaerosalinaceae” are generally greater than 16% (Fig. 3b, Table S7). In *Halobacteria* such as *Haloarcula hispanica*, *Haloferax mediterranei*, and *Natronococcus occultus* (representatives of the three order), the mole percentages of both glutamate and aspartate are increasing and are almost equal, while in “*Ca*. Nanosalinaceae” and the four “*Ca*. Nanoanaerosalinaceae” members (NHA24, NHA20, NHA23, and NHA-2, except NHA21), glutamate is unexpectedly much higher than aspartate, although both are also increasing (Fig. 3b, Table S7). The result suggests that “*Ca*. Nanosalinaceae” and most “*Ca*. Nanoanaerosalinaceae” achieved the “salt-in” with more glutamate accumulation in the proteomes, and they are different from their *Halobacteria* hosts. We infer that most “*Ca*. Nanohaloarchaeota” lineages prefer glutamate considering the two facts below: 1) AB_1215_Bin_137 contains about 12% of acidic amino acids (close to non-halophiles), and glutamate is likewise much more than aspartate (Table S7); 2) more than half of the representative genomes have glutamate dehydrogenase gene (*gdhA*) for glutamate biosynthesis (Fig. 4a, Table S8; described below). An exception is NHA21 which has almost equivalent glutamate and aspartate in its proteome (Fig. 3b, Table S7). Convergence of amino acid composition of NHA21 with *Halobacteria* may be the consequence of an excessively close connection between both. In addition, we could observe that the GC contents of most “*Ca*. Nanosalinaceae” genomes range between 39.83–46.95 mol% except for J07AB56 (with a GC content of 56.20 mol%), while those of the four “*Ca*. Nanoanaerosalinaceae” members (NHA24, NHA20, NHA23, and NHA-2) and AB_1215_Bin_137 are 38.06–41.20 mol% and 32.36 mol%, respectively (Table S2), gradually decreasing reversely along with the evolutionary process. J07AB56 was omitted because of its low-quality (Table S2). The GC content of NHA21 is up to 56.09 mol%. It may result from the further closer association with *Halobacteria* than the family “*Ca*. Nanosalinaceae”. Therefore, the hypersaline adaptation indicated by amino acid composition of proteome as well as genomic GC content also reveals the degree of intimate association with *Halobacteria* (8, 20).

**Fig. 4.**
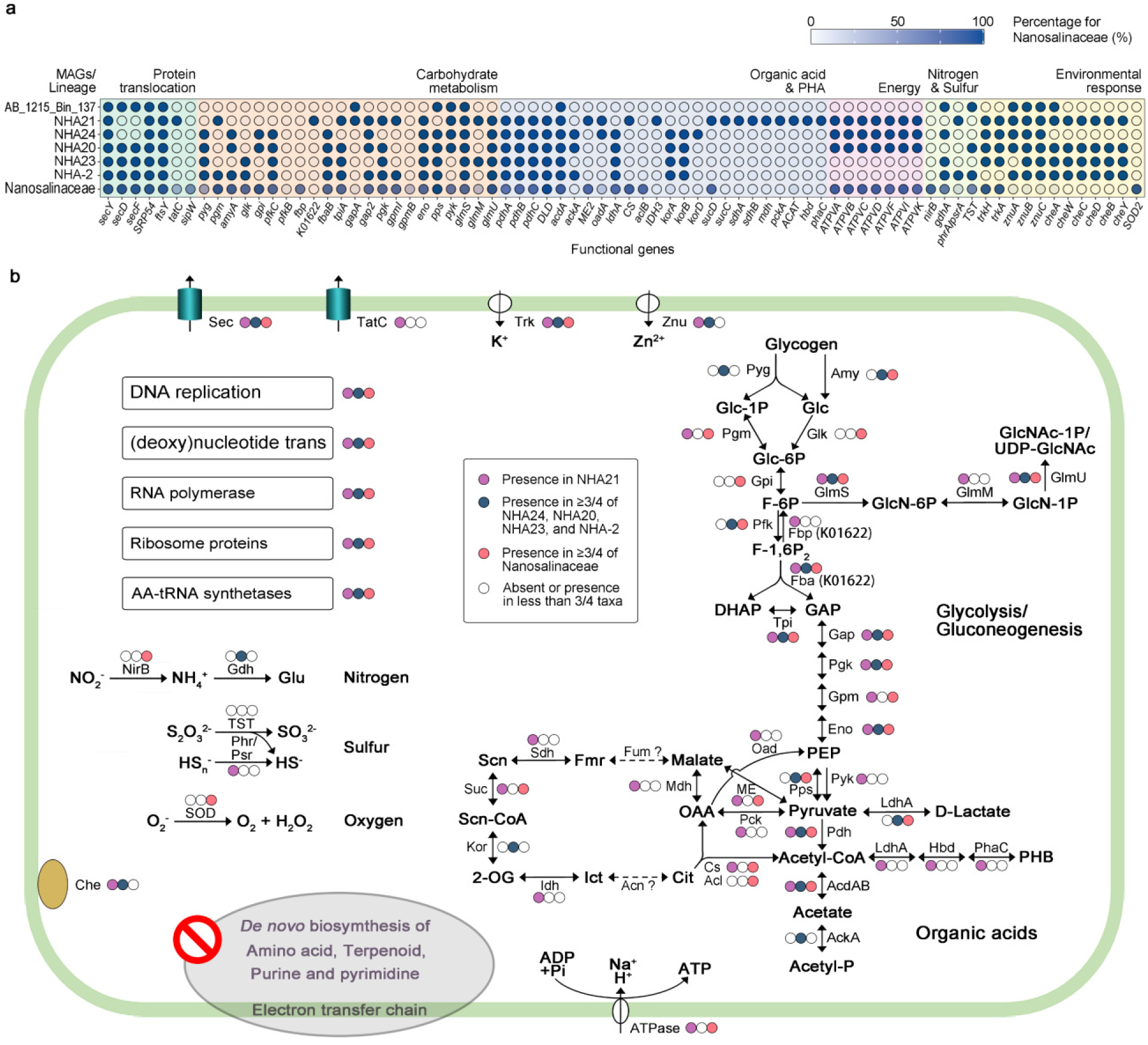
Metabolic potentials of the phylum “*Ca.* Nanohaloarchaeota” lineages. (a) Dot plot showing the presence or absence of genes involved in metabolism and environmental response in the members of “*Ca.* Nanoanaerosalinaceae” and the percentage in the family “*Ca*. Nanosalinaceae”. Solid and hollow dots indicated the presence and absence in the genomes, respectively. Transparent blues show the percentage of genes in 14 “*Ca*. Nanosalinaceae” genomes. (b) Reconstruction of metabolism potentials in three lineages of “*Ca.* Nanohaloarchaeota”. The process was estimated based on the genes involved in genetic information processing, metabolism, and environmental stress response. Solid with different colors respectively revealed the presence of the process or gene(s) in the three lineages, NHA21, the other four members of “*Ca.* Nanoanaerosalinaceae”, and “*Ca.* Nanosalinaceae”, while hollow dots indicated the absence of the process or gene(s). Glc, glucose; Glc-1P, Glucose 1-phosphate; Glc-6P, Glucose 6-phosphate; F-6P, Fructose 6-phosphate; F-1,6P_2_, Fructose 1,6-bisphosphate; DHAP, Dihydroxyacetone phosphate; G-3P, Glyceraldehyde 3-phosphate; PEP, Phosphoenolpyruvate; Glu, glutamate; OAA, oxaloacetate; Cit, citrate; Ict, isocitrate; 2-OG, 2-oxoglutarate; Scn-CoA, Succinyl-CoA; Scn, succinate; Fmr, fumarate; Che, chemotaxis; GlcN-6P, Glucosamine 6-phosphate; GlcN-1P, Glucosamine 1-phosphate; GlcNAc-1P, N-acetylglucosamine 1-phosphate; UDP-GlcNAc, UDP-N-acetylglucosamine; PHB, poly-hydroxybutyrate. Enzyme abbreviations are listed in Table S8.

Consequently, we supposed that the last common ancestor of “*Ca*. Nanosalinaceae” and “*Ca*. Nanoanaerosalinaceae” have connected with *Halobacteria*, and the two families choose different evolutionary paths. Soon, divergence appeared again in the family “*Ca*. Nanoanaerosalinaceae”. In the three lineages, “*Ca*. Nanosalinaceae” build an intimate association with *Halobacteria*, while the four “*Ca*. Nanoanaerosalinaceae” members (NHA24, NHA20, NHA23, and NHA-2) maintain a moderate connection with them. NHA21 may form a closer association with *Halobacteria*, and more evidence could not be provided until more related genomes were obtained. In a word, there are three predominantly parallel evolutionary paths found in “*Ca*. Nanohaloarchaeota”.

### “*Ca*. Nanoanaerosalinaceae” members contain divergent functional potentials

To decipher the factors that possibly drove the different evolutionary paths, the functional potentials were predicted. Generally, all genomes of “*Ca*. Nanoanaerosalinaceae” have the genes involved in the DNA replication apparatus, RNA polymerase complex, multiple ribosome proteins, and aminoacyl-tRNA biosynthesis as those of “*Ca*. Nanosalinaceae” (Table S8). Meanwhile, we found that the different genes are mainly involved in metabolism and environmental response. However, they all lacked the complete genes for the electron transfer chain and the *de novo* biosynthesis of amino acids (except glutamate), purine, pyrimidine, and terpenoid for cell membranes like “*Ca*. Nanosalinaceae” (Fig. 4, Table S8).

Nearly complete glycolysis or gluconeogenesis pathway was reconstructed in the family “*Ca*. Nanoanaerosalinaceae” (Fig. 4, Table S8). However, in the four “*Ca*. Nanoanaerosalinaceae” members NHA24, NHA20, NHA23, and NHA-2, phosphoglycerate mutase genes (*gpmI* or *gpmB*) are commonly lacking. If there was not a new gene functioning instead, this lineage might not be able to perform the glycolysis. NHA21 does not have the ADP-dependent phosphofructokinase gene (*pfkC*) like most other members in “*Ca*. Nanohaloarchaeota”, but annotates a fructose-1,6-bisphosphatase (by K01622) for gluconeogenesis. Although there is no pyruvate water dikinase gene (*pps*) in NHA21 for phosphoenolpyruvate (PEP) biosynthesis from pyruvate, its genome contains oxaloacetate decarboxylase (Na^+^ extruding; *oad*). Moreover, the glucose-6-phosphate isomerase gene (*gpi*) is lacking in NHA21, NHA23, and NHA-2; therefore, glucose-6-phosphate and fructose-6-phosphate could not be converted to each other. Some coding genes involved in *alpha*-glycan (such as glycogen) utilization were not widely present in the members of “*Ca*. Nanoanaerosalinaceae”. In summary, the two lineages of “*Ca*. Nanoanaerosalinaceae” are not so proficient in carbohydrate metabolism as “*Ca*. Nanosalinaceae”, and this may be one factor that drive the divergent symbiotic features.

Obviously, NHA21 distinguishingly harbors many genes involved in organic acids conversion (including *IDH3*, *sucC*, *sdhAB*, *mdh*, and *pckA*) and poly-hydroxybutyrate (PHB) biosynthesis (Fig. 4). Combining the incompleteness of citrate cycle in NHA21, the organic acids metabolism related genes may be involved in energy production and reducing power balance. Similarly, we also found organic acids metabolism in the other two lineages. In “*Ca*. Nanosalinaceae”, malate dehydrogenase (*ME2*), acetate-CoA ligase (ADP-forming) (*acdA*), and pyruvate dehydrogenase (*pdh*) genes are located together and even form a gene cluster in some genomes (Fig. S9a). These genes coupled the metabolism between malate, pyruvate, and acetate. Generally, pyruvate is considered a key nutrient in hypersaline environments, and it could be excreted by some members of the *Halobacteria* fed with glycerol (34). Meanwhile, glycerol is an important osmotic stabilizer produced by *Dunaliella*, one leading primary producers in hypersaline ecosystems (35). Therefore, pyruvate might be consumed with acetate and malate as products. In this process, ATP was generated, and reducing power was balanced (Fig. S9b). In the lineage of the four “*Ca*. Nanoanaerosalinaceae” members (NHA24, NHA20, NHA23, and NHA-2), *ME2* gene is lacking, and other genes are not linked (Table S8). However, they commonly have lactate dehydrogenase gene (*ldhA*), whose protein product could replace the role of ME2 in NAD^+^ regeneration (Fig. S9c). Meanwhile, their secretions are also different. Hence, it is inferred that the metabolic variances of organic acids in different lineages may affect the degree of intimacy with different *Halobacteria*.

Likewise, the different metabolic potentials such as the nitrite reduction, ammonia assimilation, thiosulfate and polysulfide reduction (Fig. 4) may also interfere in the symbiosis with *Halobacteria*.

In addition to metabolism, some environmental responses are different in the three lineages. The subunits SecYDF of the Sec-dependent protein export system are present in almost all of the 20 representative genomes. We found the TatC subunit of the twin-arginine translocation system in NHA21, as well as some of the family “*Ca*. Nanosalinaceae”, but not in the lineage of the other four “*Ca*. Nanoanaerosalinaceae” members (Fig. 4a, Table S8). The Tat system exports the folded protein, and this process can avoid the protein denaturation during the unfolding and refolding under the hypersaline conditions (36). Consistently, the TatC prefers to exist in the lineages with acidic proteome (imply high intracellular salinity). Furthermore, we observed that chemotaxis related genes (*cheAWCDBY*) and zinc transporter genes (*znuABC*) are present in “*Ca*. Nanoanaerosalinaceae”, while they are not in any of the 14 representative genomes of “*Ca*. Nanosalinaceae” (Fig. 4a). It was reported that zinc was important for bacterial chemotaxis (37), and it may play a similar role in “*Ca*. Nanoanaerosalinaceae”. Furthermore, “*Ca*. Nanoanaerosalinaceae” survive in hypersaline sediment (Table 1), and the sediment is not so inhabited by *Halobacteria* as brine (28, 38). The chemotaxis is possibly significant to seek out the *Halobacteria* hosts. Conversely, an Fe-Mn family superoxide dismutase gene (*SOD2*) is only annotated in the family “*Ca*. Nanosalinaceae” but not in “*Ca*. Nanoanaerosalinaceae” (Fig. 4, Table S8). Accordingly, superoxide dismutase may play a part in response to the reactive oxygen species superoxide in aerobic environments. Equally, these environmental response genes may also be the factors for the divergent evolution.

### Inference of historical events driving the evolution

In order to interpret the evolutionary paths in “*Ca*. Nanohaloarchaeota” including “*Ca*. Nanoanaerosalinaceae”, the historical events are decoded during the speciation of three lineages in “*Ca*. Nanohaloarchaeota” over the functional potentials of each member. Firstly, we identified homology relationships between the 32,540 CDSs from the 20 “*Ca*. Nanohaloarchaeota” as well as 9 EX4484-52 genomes. In total, we obtained 3,007 orthogroups, and 80.6% of the CDSs are assigned to one of the orthogroups. For each of 20 “*Ca*. Nanohaloarchaeota” genomes, at least 66.1% of CDSs are included in these orthogroups. In the five representative genomes of “*Ca*. Nanoanaerosalinaceae”, 66.1% of 1285 CDSs in NHA21, and 92.2–94.6% of CDSs in the other four genomes are included. So, the orthogroups could reflect the dominant functions in the “*Ca*. Nanoanaerosalinaceae” branch. Afterward, we performed the clustering analysis of functional profiles based on all the 3,007 orthogroups using the non-metric multidimensional scaling (NMDS) method. The result revealed that the fourteen “*Ca*. Nanosalinaceae” and nine EX4484-52 members gather together, respectively (Fig. S10). AB_1215_Bin_137 is close to one EX4484-52 member. Meanwhile, four “*Ca*. Nanoanaerosalinaceae” members NHA24, NHA20, NHA23, and NHA-2 form another cluster, while NHA21 is far away from this cluster. The NMDS result agrees with the HGT, hypersaline adaptation, and functional analyses that the two families could be classified into three lineages with different evolutionary paths.

Subsequently, ancestor genomes and historical events (originations, duplications, transfers, and losses) were approximately based on the orthogroups using the Amalgamated likelihood estimation (ALE) approach (39, 40). Briefly, 1,629 orthogroups containing 4 or more sequences were selected. The sequences of each orthogroup were aligned, trimmed, and then used to reconstruct the UFBOOT gene trees. Among them, 124 single-copy orthogroups distributed in no less than 18 “*Ca*. Nanohaloarchaeota” genomes and no less than 5 EX4484-52 genomes were selected to rebuild a consensus ultrafast bootstrap approximation result as a species tree. The unrooted tree reconstructed using IQ-TREE was restructured by setting the tree root (employing the midpoint method). The root was located at the node between the clade of “*Ca*. Nanohaloarchaeota” and EX4484-52. The tree structure of the three lineages was in line with the other three phylogenetic analyses based on 122 archaeal single-copy conserved proteins of GTDB, ribosome proteins, and 16S rRNA gene (Figs. S4–S6).

ALE was performed based on the consensus species tree and 1,629 genes’ UFBOOT trees to estimate the frequencies of ancestral events and copies. Then the outputs were parsed using the scripts provided in the previous literature (41). The output named DTLO table was obtained, and the data on the frequencies of historical events at each node or the last common ancestor (LCA) of the subordinate taxa were exhibited in the “*Ca*. Nanohaloarchaeota” tree with EX4484-52 as an outgroup (Fig. 5a). The result revealed that the inferred genome sizes (copy number frequency in DTLO table) sharply increase from node 54 (the LCA of “*Ca*. Nanosalinaceae” and “*Ca*. Nanoanaerosalinaceae”) to node 52 (the LCA of “*Ca*. Nanoanaerosalinaceae”), then node 50 and node 46 (Fig. 5a). In addition, we could observe a similar trend in “*Ca*. Nanosalinaceae” branch, such as from node 54 to node 53, node 51, and then node 49. Correspondingly, there are high frequencies of gene originations and transfers. In fact, the gene originations are composed of transfers from the lineages outside the tree and are true originations (42), while the gene transfers are from the lineages inside the tree. Moreover, the frequencies of gene losses are high at node 50 and node 53, while those of gene duplications are generally low. This result illustrated that the historical events, especially the acquisition of genes occurred before or at the beginning of family emergence.

**Fig. 5.**
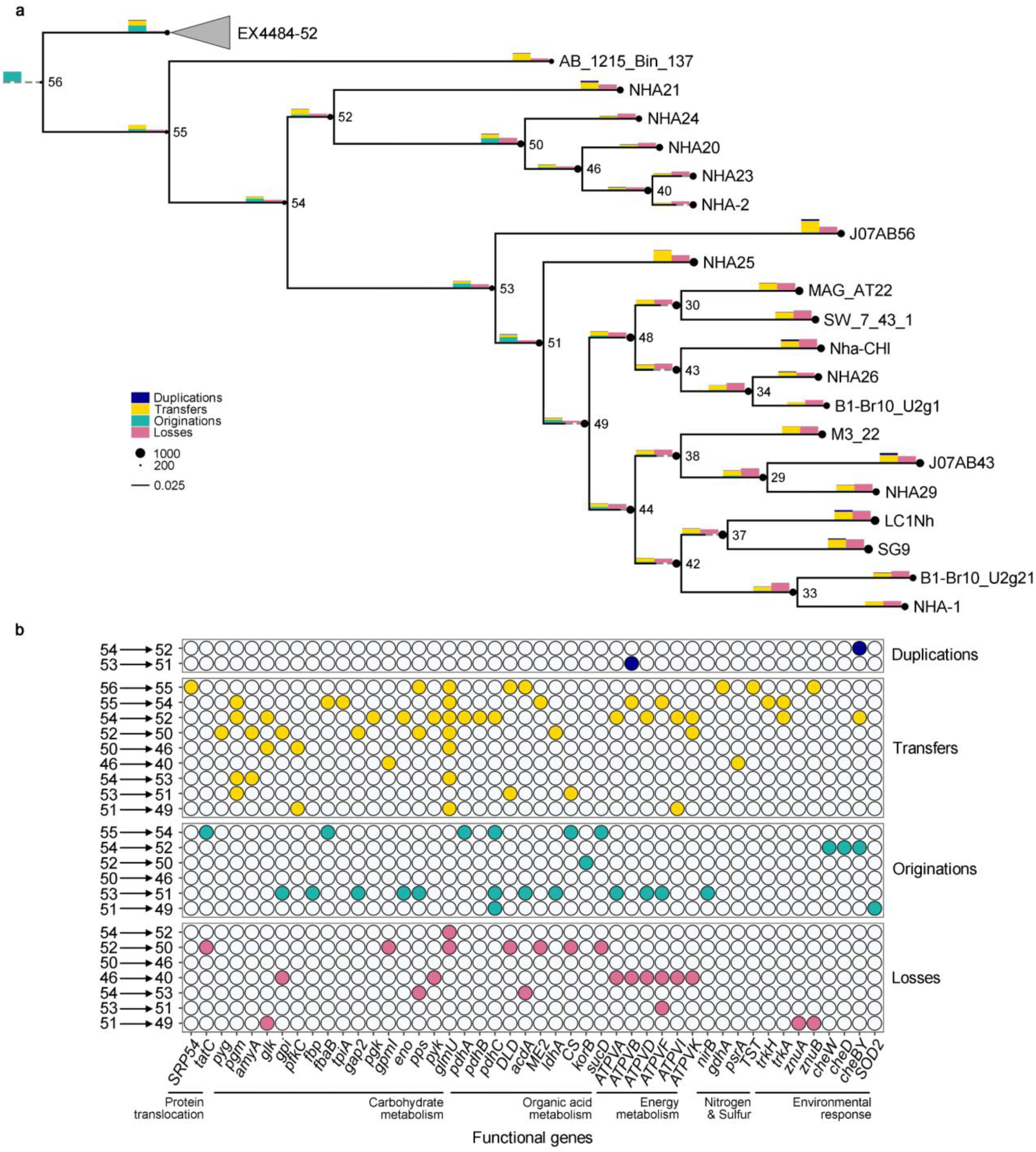
History events approximation in the phylum “*Ca*. Nanohaloarchaeota”. (a) Ancestral reconstruction tree of “*Ca*. Nanohaloarchaeota”. The consensus tree of ultrafast bootstrap approximation based on 124 single-copy ubiquitous orthogroups in representative genomes is exhibited. The historical events are approximated based on the species tree and 1,629 gene UFBOOT trees. The black circles at nodes are their (inferred) genome sizes represented by the radius. Some internal nodes of interest are marked by Arabic numerals. The bar charts above the horizontal branches represent the numbers of duplications, transfers, origination, and losses (bar heights in legend correspond to 300 events each). Some branches were extended with dashed lines to fit the width of the bar charts. The branches leading to the phylum EX4484-52 were collapsed. (b) Dot plot showing the functional annotation of the historical event at the interest nodes. The orthogroups that achieved a threshold of 0.3 in the raw reconciliation frequencies were reported. The putative functions of orthogroups are estimated by medoid sequences which have the highest sum of similarity scores with all other sequences based on the BLOSUM62 substitution matrix.

To find out the putative functions involved in the historical events, we selected the orthogroups that achieved a threshold of 0.3 in the raw reconciliation frequencies to avoid misses of true events (42). The results revealed that many genes of glycolysis or gluconeogenesis, including *gpi*, *fbp*, *gap2*, *eno*, and *pps* are originations from node 53 to node 51 (Fig. 5ab). Meanwhile, some genes encoding V/A-type H^+^/Na^+^-transporting ATPase subunits A, D, F, and nitrite reductase (NADH) large subunit gene *nirB* are also inferred for the originations at this node. The ATP synthase of “*Ca*. Nanohaloarchaeota” are reported to share a sisterhood relationship with *Halobacteria* (21). Next, we focused on the “*Ca*. Nanoanaerosalinaceae” branch and its latest ancestor. We found that pyruvate dehydrogenase genes *pdhAC* and some organic acids metabolism related genes are originations from node 55 to node 54. Especially pyruvate dehydrogenase is widely present in the three lineages, and it is seemingly indispensable. In addition, *tatC* is also origination from node 55 to node 54, and then it was lost from node 52 to node 50 (Fig. 5ab). Factually, it is impossible to infer the historical events in NHA21, and two reasons are: 1) many genes (about one third) are not included in the orthogroups, and 2) there is only one genome in this lineage and the specific genes would not form orthogroups with four or more sequences to perform the ALE analysis. However, the Tat protein translocation system was obtained by the LCA of the three lineages, and it is lost by the four “*Ca*. Nanoanaerosalinaceae” members (NHA24, NHA20, NHA23, and NHA-2). The genes like *tatC* in NHA21 could be clarified, including *ME2* and *CS* genes (Fig. 5ab).

The result also exhibited that the chemotaxis related genes *cheWDY* are originations from node 54 to node 52, while the *SOD2* gene is origination from node 51 to node 49 (Fig. 5ab). These genes were found present in the subordinate genomes (Fig. 4a) described above, and they play important roles in the environmental response and symbiosis with *Halobacteria*. Consequently, the inferred historical events enrich the evolutionary details of three lineages and further confirm the divergent lineages in “*Ca*. Nanohaloarchaeota”.

## Conclusion

We report a novel family “*Ca*. Nanoanaerosalinaceae” represented by five de-replicated genomes. This family is placed at the root of the phylogenetic trees of the phylum “*Ca*. Nanohaloarchaeota”. Combining the hypersaline adaptation and the proportion of horizontally acquired genes from *Halobacteria*, we deduced that the family arose at the early stage of the development of *Halobacteria*-associated symbiosis. Meanwhile, we classify the family “*Ca*. Nanoanaerosalinaceae” into two separate lineages by following their divergence from the family “*Ca*. Nanosalinaceae”, the third lineage. By predicting the functional potentials and inferring the historical events, we illuminate the primarily evolutionary process of many lineage specific genes involved in carbohydrate metabolism, organic acids metabolism, chemotaxis, reactive oxygen species response (*SOD2*), and hypersaline adaptation (Tat protein translocation system). The family “*Ca*. Nanoanaerosalinaceae” broaden the archaeal diversity of “*Ca*. Nanohaloarchaeota”. This research sheds light on the evolution of symbiosis with *Halobacteria*, and will provide a model for similar studies in other DPANN lineages.

### Description of the family **“***Ca*. Nanoanaerosalinaceae**”** and taxa classified in the family

Description of “*Ca*. Nanoanaerosalinaceae” fam. nov. (Na.no.an.ae.ro.sa.li.na.ce’ae, N.L. fem. n. *Nanoanaerosalina*, Candidatus generic name; -*aceae*, ending to designate a family; N.L. fem. pl. n. *Nanoanaerosalinaceae*, the *Nanoanaerosalina* family). The type genus of the family is the genus “*Candidatus* Nanoanaerosalina”.

Description of “*Ca*. Nanoanaerosalina” gen. nov. (Na.no.an.ae.ro.sa.li’na. Gr. masc. n. *nanos*, a dwarf; Gr. pref. *an*-, not; Gr. masc. n. *aêr* (*gen*. *aeros*), air; N.L. masc. adj. *salinus*, saline; N.L. fem. n. *Nanoanaerosalina*, a dwarf saline organism not living in air). The type species of the genus is “*Candidatus* Nanoanaerosalina halalkaliphila”.

Description of “*Ca*. Nanoanaerosalina halalkaliphila” sp. nov. (hal.al.ka.li’phi.la. Gr. masc. n. *hals*, (*gen*. *halos*), salt; N.L. neut. n. *alkali*, alkali; N.L. adj. *philus -a -um*, friend, loving; from Gr. adj. *philos -ê -on*, loving; N.L. fem. adj. *halalkaliphila*, salt and alkali-loving). The type material is the metagenome-assembled genome NHA20, whose GenBank accession number (WGS) is JALDAE000000000.

## Materials and Methods

### Enrichment culture and metagenomic sequencing

Deep sediment samples from five crystallizer ponds of different salinities (1, 3, 15, 24, and 33%) described in the previous study (28) were used for enrichment culture. In brief, 5 g of mixed four sediment samples (from the identical pond) was added with 5 mL of corresponding brine and 5 mL sterile medium with the same salinity as the brine. The five media were composed of (per liter): 0.2 mg MgCl_2_•6H_2_O, 0.05 g KH_2_PO_4_, 2 g KCl, and variable concentrations of NaCl, NaHCO_3_, and CaCl_2_ for samples from different ponds.

Their concentrations were: 10 g, 2 g, and 1.2 mg, respectively, for the 1% samples; 30 g, 3 g, and 0.4 mg for 3%; 150 g, 13 g, and 6.5 mg for 15%; 240 g, 28 g, and 7.7 mg for 24%; 370 g, 5 g, and 4.1 mg for 33%. After sterilization, pH values usually approximate 10.0. After premixing the samples with brine and media, a final concentration of 2.42 g/L Na_2_MoO_4_•2H_2_O, 5 mg/L kanamycin, 20 mg/L ampicillin, 2.4 g/L NaS_2_•9H_2_O, and then 0.52 g/L sodium formate dihydrate (signified “F” in sample), 0.68 g/L sodium acetate trihydrate (signified “A” in sample) or no substrate was added (details shown in Table S1). A total of fifteen samples were anaerobically and statically incubated at 30 °C without light for about 210 days. Then, microbial cells with insoluble matter were collected using centrifugation at 4 °C. Total DNA was extracted from the samples using a PowerSoil DNA Isolation kit (MoBio, CA, United States) for metagenome sequencing. However, DNA was not enough from the sample of 1% salinity with no substrate. So, fourteen competent DNA samples were used for subsequent library construction and metagenomic sequencing in the Illumina platform to generate 150 bp paired end reads (Table S1).

### Contig assembly and genome binning

Quality control of raw reads of each metagenome was performed using the read_qc module with default parameters of metaWRAP v1.2.2 pipeline (43). Clean reads generated for each sample were individually assembled into contigs using the assembly module of metaWRAP with default assembler MEGAHIT v1.1.3 (44), and short contigs (<1,000 bp) were removed. Three different metagenomic tools for genome binning, CONCOCT v1.0.0 (45), MetaBAT2 v2.12.1 (46), and MaxBin2 v2.2.6 (47) integrated into the binning module of metaWRAP, were used to recover initial MAGs from each metagenome, respectively. In addition, to obtain more “*Ca*. Nanohaloarchaeota” MAGs, eighteen metagenomes of brine and surface sediment samples from our previous study (23) were re-analyzed by following the genome binning approach as described below.

Then, all MAGs obtained from the same metagenome were individually refined with a minimum completement of 50% or 30% and a maximum contamination of 10% or 20% using the bin_refinement module in metaWRAP. Subsequently, the best representative genomes were chosen from all the refined MAGs using dRep v3.2.0 with a threshold of 99% ANI (48).

### Genomes collection, quality estimation, and functions prediction

The genome sequences of “*Ca*. Nanohaloarchaeota” and its closely related lineages, including the phyla “*Ca*. Aenigmarchaeota”, EX4484-52, PWEA01, and QMZS01, were collected (Table S2) by October 9, 2021. Firstly, 232 genomes were downloaded from the GenBank of the National Center for Biotechnology Information (NCBI, https://www.ncbi.nlm.nih.gov/) under the following taxonomy IDs: 743724 for the phylum “*Ca*. Aenigmarchaeota”, 1462430 for the phylum “*Ca*. Nanohaloarchaeota” in the DPANN group, 1051663 for the class “Nanohaloarchaea” in the phylum *Euryarchaeota*, 2565780 for the unclassified DPANN group. Two genomes of “*Candidatus* Nanohaloarchaeum antarcticus” (16) were obtained via the identification numbers 2643221421 and 2791354821 in the Integrated Microbial Genomes (IMG, https://img.jgi.doe.gov/). Additionally, 148 archaeal genomes of our previous studies (23, 28) were also used. Besides, 2,285 archaeal representative genomes in GTDB were downloaded (Table S3) for phylogeny analysis.

Taxonomy was generally pre-analyzed using classify workflow in GTDB-Tk (V1.7.0) (49) based on Release 202 in Genome Taxonomy Database (https://gtdb.ecogenomic.org/).

Subsequently, basic statistics (including contigs number, genome size, N50, N90, and GC content) of genomes were generated using bbstats.sh script (Last modified July 25, 2019) in BBTools suite (sourceforge.net/projects/bbmap/). The genomes were annotated using Prokka (version 1.13) with the settings of Archaea for annotation mode and RNAmmer (v1.2) for rRNA prediction (50, 51). In this process, Prodigal (v2.6.3) was used to find protein-coding features (52). The number of genes, CDSs, rRNAs, and tRNA were retrieved from the Prokka output. Genome completeness and contamination were estimated using lineage-specific workflow in CheckM (v1.1.3) (53). The isoelectric point of each protein was predicted using the “Protein isoelectric point calculator” (54). Functions of CDSs in each genome were predicted using the diamond method in eggNOG-mapper (v2.0.0) (55, 56). From the output, Clusters of Orthologous Genes (COG) categories (57) and KEGG (stands for Kyoto Encyclopedia of Genes and Genomes) Orthology (KO) identifiers were retrieved. Metabolic pathways were reconstructed using the online tool KEGG Mapper with KO annotation (https://www.genome.jp/kegg/). Carbohydrate-active enzymes were annotated using dbCAN2 (v2.0.6) based on the CAZy database version CAZyDB.07312019 (58, 59).

### Phylogeny and sequence identity analyses

The phylogenetic trees were based on 122 archaeal ubiquitous single-copy proteins, ribosome proteins, and 16S rRNA gene with 2,285 archaeal genomes except NATU lineages in GTDB as an outgroup. Briefly, the multiple sequence alignment file of the 122 archaeal ubiquitous single-copy proteins were also produced using the classify workflow in GTDB-Tk (49). The consensus tree of ultrafast bootstrap approximation was reconstructed using IQ-TREE (multicore version 1.6.12) with an ultrafast bootstrap of 1,000 and standard model selection followed by tree inference (60). The best-fit model was chosen according to Bayesian Information Criterion. The archaeal ribosome proteins were inferred from the genomes and then aligned, masked, and trimmed using AMPHORA2 with default options (61). The aligned and trimmed amino acid sequences of Rpl2p, Rpl3p, Rpl4lp, Rpl5p, Rpl6p, Rpl14p, Rpl18p, Rpl22p, Rpl24p, Rps3p, Rps8p, Rps10p, Rps17p, and Rps19p were concatenated using AMAS (62). The phylogenomic tree was reconstructed using IQ-TREE with the same settings, and the best-fit model was chosen. The archaeal 16S rRNA gene sequences were predicted using RNAmmer (v1.2) (51). Those of 1,450–1,700 bp length and high-quality (more than 1100 score and no ambiguous sites) were selected (Table S4), referring to the previous pieces of literature (51, 63). The sequences were aligned using MUSCLE v3.8.31 with default options (64). The phylogenetic reconstruction was also done using the same method, and the best-fit model was chosen. The phylogenetic trees were visualized using the Interactive Tree of Life (iTOL, version 6.5) (65). The archaeal unrooted life trees were generally reconstructed by setting the root at the node between the DPANN group and other lineages. The phylogenetic trees of NATU clade were reconstructed by following the same approach.

The identity between two 16S rRNA gene sequences was estimated using Nucleotide-Nucleotide BLAST 2.6.0+ (66). Average amino acid identity (AAI) between two predicted proteomes was calculated using the online AAI calculator (http://enve-omics.ce.gatech.edu/aai/index). Average Nucleotide Identity (ANI) between two genomes was computed using FastANI (version 1.33) with default options (67).

### Horizontal gene transfer events prediction

The putative horizontal gene transfer (HGT) events were computed using the HGTector pipeline version 2.0b3 (32). Firstly, the GTDB taxonomy-based database for HGTector analysis was built by considering that different genomes of “*Ca*. Nanohaloarchaeota” were assigned to *Euryarchaeota* or DPANN group. In brief, the taxonomy file was downloaded from GTDB within Release 202 data, and then it was reformatted into NCBI taxdump style using a Python 3 script named gtdb_to_taxdump.py provided by HGTector contributors (https://github.com/qiyunlab/HGTector). Three files named names.dmp, nodes.dmp and taxid.map were produced. All proteins of 2,339 archaeal representatives in GTDB were directly downloaded from GenBank or predicted based on the genomes using Prodigal (v2.6.3) (52). GTDB-based and local prot.accession2taxid file was created based on the produced taxid.map file and all proteins. The database was built from all the proteins and the GTDB-based taxonomy files using makedb command in DIAMOND v0.9.26.127 (56). After that, a batch homology search was performed using a diamond method for each proteome. HGT events were predicted using the analyze command with the following settings: “self” group, “*Ca*. Nanohaloarchaeota” (whose taxid in the local GTDB-based taxonomy was 18); “close” group, “*Ca*. Nanohaloarchaeota” and EX8848-52 (taxid: 20); maximum number of hits, 12; maximum E-value cutoff, 1e-8; minimum percent identity cutoff, 30%; minimum percent query coverage cut off, 50%; bandwidth for Gaussian KDE, auto. Donor’s taxonomy of HGT-derived gene was deciphered using lineage command in TaxonKit (v0.9.0) (68).

### Comparative genomic analysis for ancestral reconstruction

The comparative genomic analysis was performed by referring to the published research (41, 42, 69). Briefly, orthogroups were found from the genome set (20 “*Ca*. Nanohaloarchaeota” and 9 EX4484-52 genomes) using OrthoFinder version 2.4.0 with default settings (70). The gene counts matrix of orthogroups in the genomes was used to compare the functional profile. Non-metric Multidimensional Scaling (NMDS) analysis was performed using the metaMDS function with default options in R package vegan v2.5-7 (https://github.com/vegandevs/vegan/).

We retained those orthogroups with 4 or more genes according to the previous literature (69). The genes of identical orthogroup were aligned using MAFFT v7.407 with L-INS-i method of high accuracy (71). The columns were removed using the heuristic automated1 method of trimAl version 1.2rev59 (72), and then the sequences containing too many gaps were abandoned with the following options: minimum overlap of a position with other positions, 0.3; minimum percentage of the satisfied positions, 50. For the reconstruction of the species tree, 124 single-copy orthogroups were manually selected that were present in no less than 18 “*Ca*. Nanohaloarchaeota” genomes (≥90%) and no less than 5 EX4484-52 genomes (>50%). The aligned and trimmed sequences were concatenated using AMAS (62). Correspondingly, the phylogenomic tree was reconstructed using IQ-TREE, and the best-fit model of LG+F+I+G4 was chosen. The root of this species tree was reset using the midpoint method, a built-in function in iTOL (65). To obtain the UFBOOT trees of 1,629 orthogroups, we used IQ-TREE with the same settings (-m, LG+G; -bb, 1000; -wbtl) as in the previous research (69). The frequencies of duplications, transfers (gene transfers from the lineage inside the species tree), losses, and originations (gene transfers from the lineages outside the species tree, or true gene originations), as well as copy numbers of 1,629 orthogroups at each node, were inferred using the way of maximum likelihood estimation (ALEml_undated command) in ALE v0.4 (39). The number of each event and genome size were inferred by parsing the “.uml_rec” files of ALE output using the Python scripts set named ALE helper (41). Ancestral reconstruction tree was visualized using ETE Toolkit version 3.1.2 (73).

The orthogroups that achieved a threshold of 0.3 in the raw reconciliation frequencies were counted. This threshold is relaxed but necessary to avoid misses of many true events (42). A medoid sequence was selected from each orthogroup as the representative for functional annotation. The medoid sequences have the highest sum of similarity scores with all other sequences based on the BLOSUM62 substitution matrix using Protein-Protein BLAST 2.6.0+ (66).

## Data availability

The “*Ca*. Nanohaloarchaeota” genomes are available from the NCBI under the BioProject identifier PRJNA797678. DNA sequencing data have been deposited in BioProject identifiers PRJNA549802 and PRJNA679647. Metagenomic sequencing data of 14 enrichment samples are deposited in BioProject identifiers PRJNA769545. Raw data (including protein files, tree files, horizontal gene transfer analysis, comparative genomics files, etc.) generated in this study are available through [https://doi.org/10.6084/m9.figshare.19549495].

## Acknowledgements

This study was funded by the National Natural Science Foundation of China (No. 91751201, 32000046), and partially supported by the Senior User Project of RV KEXUE (No. KEXUE2019GZ05) provided by the Center for Ocean Mega-Science, Chinese Academy of Sciences.

The authors declare no competing interests.

